# Ester-linked ubiquitin sites formed by HOIL-1 in TLR7 signaling include four novel ubiquitin dimers

**DOI:** 10.1101/2022.09.28.509890

**Authors:** Elisha H. McCrory, Vyacheslav Akimov, Philip Cohen, Blagoy Blagoev

**Affiliations:** MRC Protein Phosphorylation and Ubiquitylation Unit, University of Dundee, Scotland, UK; University of Southern Denmark, Odense, Denmark

**Keywords:** Ubiquitylation, HOIL-1, E3 ligase, Toll-Like Receptors, Innate immunity

## Abstract

The E3 ligase HOIL-1 forms ester bonds *in vitro* between ubiquitin and serine/threonine residues in proteins. Here, we exploit UbiSite technology to identify serine and threonine residues undergoing HOIL-1 catalysed ubiquitylation in macrophages stimulated with R848, an activator of the TLR7/8 heterodimer. We identify Thr12, Thr14, Ser20 and Thr22 of ubiquitin as amino acid residues forming ester bonds with the C-terminal carboxylate of another ubiquitin molecule, increasing from 8 to 12 the different types of ubiquitin dimer formed in cells. We also identify Ser175 of IRAK4, Ser136, Thr163 and Ser168 of IRAK2 and Thr141 of MyD88 as further sites of HOIL-1-catalysed ubiquitylation together with lysine residues in these proteins that also undergo R848-dependent ubiquitylation. These findings establish that the ubiquitin chains attached to components of myddosomes are initiated by both ester and isopeptide bonds. Ester bond formation takes place within the proline, serine, threonine-rich (PST) domains of IRAK2 and IRAK4 and the intermediate domain of MyD88. The ubiquitin molecules attached to Lys162, Thr163 and Ser168 of IRAK2 are attached to different IRAK2 molecules.

## Introduction

The Linear Ubiquitin Assembly Complex (LUBAC) contains two different E3 ubiquitin ligases, HOIL-1 (haem-oxidised IRP2 ubiquitin ligase-1), also called RBCK1 (RING-B-Box-coiled-coil protein interacting with PKC 1), and HOIP (HOIL-1-interacting protein). HOIP catalyses the formation of Met1-linked ubiquitin (M1-Ub, also called linear ubiquitin) in which the α-amino group of the N-terminal methionine residue of ubiquitin forms a peptide bond with the C-terminal carboxylate of another ubiquitin molecule **(Kirisako *et al*, 2006)**. In contrast, HOIL-1 is one of the few E3 ligases that forms ester bonds between the C-terminal carboxylate of ubiquitin and serine and threonine residues in proteins **(Kelsall *et al*, 2019; Rodriguez Carvajal *et al*, 2021)**, the great majority of E3 ligases forming isopeptide bonds between the C-terminus of ubiquitin and ε-amino side chains of lysine residues, including the seven lysine residues in ubiquitin **(Vijay-Kumar *et al*, 1987)**.

One well established physiological role of LUBAC is to regulate innate immune signaling pathways in which M1-Ub oligomers formed by HOIP interact with NEMO (NF-κB Essential Modulator) **(Lo *et al*, 2009; Rahighi *et al*, 2009)**, a component of the canonical IκB kinase (IKK) complex. This induces a conformational change that permits the TAK1 kinase complex to initiate IKK activation **(Zhang *et al*, 2014)** leading to activation of the transcription factors NF-κB and IRF5 (Interferon Regulatory Factor 5) **(Lopez-Pelaez *et al*, 2014; Ren *et al*, 2014)** that are essential for production of the inflammatory mediators needed to mount responses that combat infection by microbial pathogens. The M1-Ub chains also interact with A20 **(Bosanac *et al*, 2010; Skaug *et al*, 2011)** and ABIN1 (A20-binding inhibitor of NF-κB 1), which function to restrict TAK1 and IKK activation, preventing the overproduction of inflammatory mediators that cause inflammatory and autoimmune diseases **(Nanda *et al*, 2019; Nanda *et al*, 2011)**. The M1-Ub oligomers formed by HOIP therefore have both positive and negative roles in controlling inflammatory mediator production.

The HOIL-1 E3 ligase also appears to have positive and negative roles in the regulation of immunity. Macrophages in which wild type HOIL-1 is replaced by the E3 ligase inactive HOIL-1[C458S] mutant produce reduced amounts of the pro-inflammatory cytokines IL-6 and IL-12 in response to TLR (Toll-Like Receptor)-activating ligands, whereas cytotoxic T cells from the same mice produce enhanced amounts of interferon γ and GM-CSF (Granulocyte Macrophage Colony Stimulating Factor) in response to IL-18 **(Petrova *et al*, 2021)**. HOIL-1-catalysed ester-linked ubiquitylation events can therefore decrease or increase inflammatory mediator production, depending on the ligand, pathway, and cell type. This is consistent with the observation that HOIL-1-deficient humans display a combination of immunodeficiency and auto-inflammation **(Boisson *et al*, 2012; Phadke *et al*, 2020)**. However, the molecular mechanisms by which ester-linked ubiquitylation regulates immunity are unknown.

IL-18, other IL-1 family members and TLRs all initiate intracellular signaling by forming myddosomes **(Lin *et al*, 2010; Motshwene *et al*, 2009)**, which are large oligomeric complexes comprising MyD88 (myeloid differentiation primary response gene 88), IRAK4 (interleukin receptor activated kinase 4) and IRAKs 1 and 2. All four components undergo polyubiquitylation during IL-1 or TLR signaling, the ubiquitin oligomers attached to these proteins containing both Lys63-ubiquitin (K63-Ub) linkages formed by the E3 ligases TRAF6, Pellino 1 and Pellino 2 **(Deng *et al*, 2000; Ordureau *et al*, 2008; Strickson *et al*, 2017)** and M1-Ub linkages formed by HOIP **(Emmerich *et al*, 2013)**. Some, but not all these hybrid ubiquitin chains, appear to be initiated by the formation of HOIL-1-catalysed ester bonds, because they are cleaved at mildly alkaline pH by the nucleophile hydroxylamine, and the hydroxylamine-sensitive bonds are not formed in macrophages from HOIL-1[C458S] mice (**Kelsall *et al***., **2019**). However, whether these inferences are correct, requires the specific amino acid residues to which the ubiquitin chains are attached to be identified. Here, we have used “UbiSite” technology to conduct this analysis as it enables sites of ester-linked and isopeptide-linked ubiquitylation to be identified in the same experiment. These studies have also revealed that HOIL-1 forms ester-bonds between two different ubiquitin molecules during TLR signaling.

## Results and Discussion

### Some of the ubiquitin chains attached to IRAK4 during R848 signaling are cleaved by hydroxylamine

Although IRAK4 undergoes polyubiquitylation during TLR ligation **(Emmerich *et al***., **2013)**, whether the ubiquitin chains attached to this protein are initiated by the formation of isopeptide or ester bonds has not been investigated. Here we found that, as for IRAK1 and IRAK2 (**lower panels of Figs 1A and 1B**), some of the ubiquitylated IRAK4 formed after stimulation for 20 min or 6 h with R848 (**upper panels of Figs 1A and 1B**) are converted to unmodified IRAK4 by treatment with hydroxylamine. Thus, some of the ubiquitin chains attached to IRAK4 appear to be initiated by ester-bond formation.

**Figure 1:**
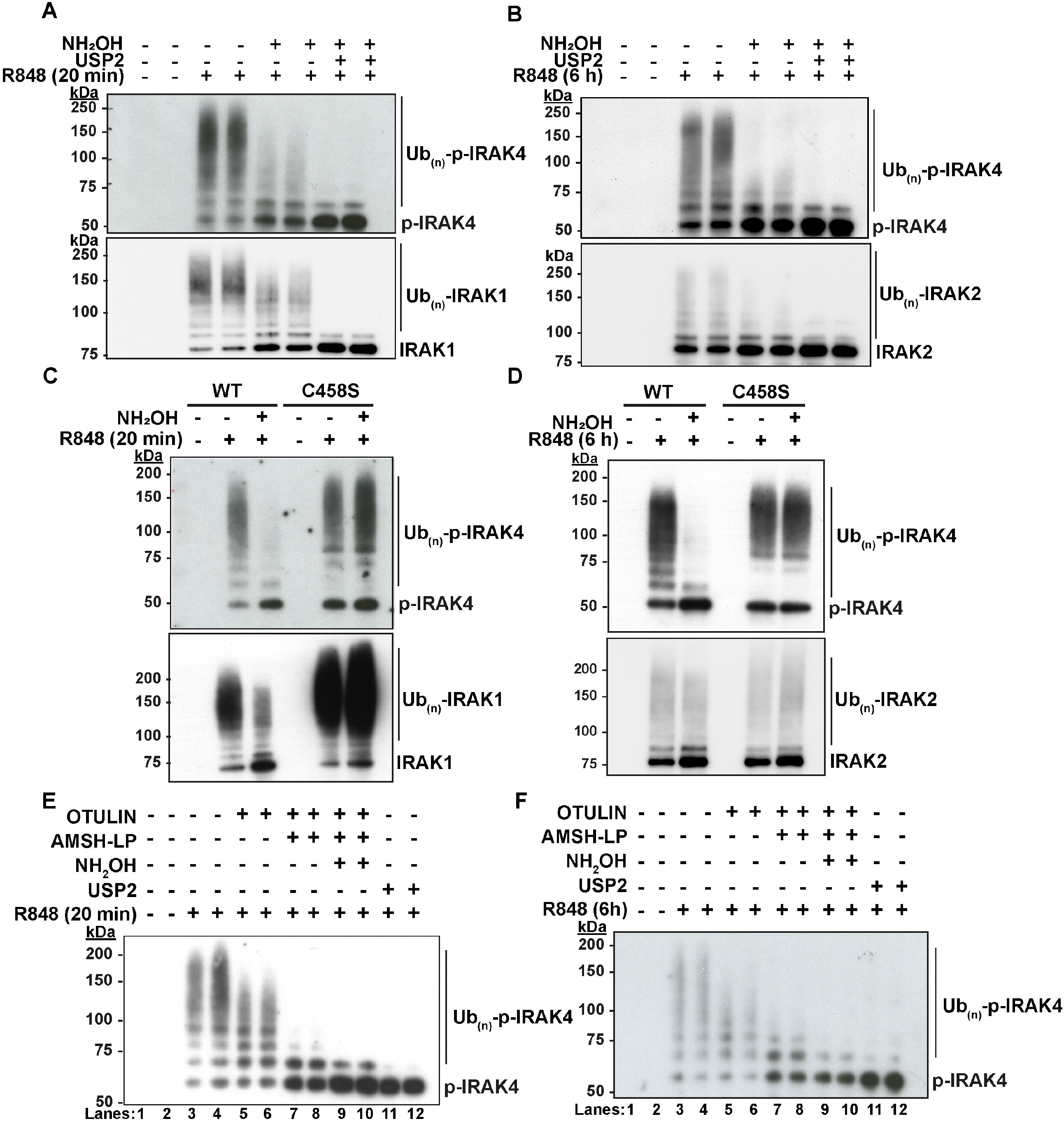
Some of the ubiquitin chains attached to IRAK4 in R848-stimulated macrophages are initiated by ester bond formation catalysed by HOIL-1. A, B. RAW264.7 cells were stimulated for 20 min (A) or 6 h (B) with the TLR7-activating ligand R848 (250 ng/ml) and ubiquitylated proteins were captured from the cell extracts on Halo-NEMO beads. The Halo NEMO pull downs were incubated for 1 h without (−) or with (+) 0.5 M hydroxylamine at pH 9.0 followed by incubation without (-) or with (+) the broad specificity deubiquitylase USP2 (1 μM). C,D. The ubiquitylated proteins captured from the extracts of BMDM from wild type (WT) or HOIL-1[C458S] mice (C458S) were incubated for 1 h without (−) or with (+) 0.5 M hydroxylamine at pH 9.0 and analysed as in A, B. E,F. As in A, B except that, prior to SDS-PAGE and immunoblotting, the Halo-NEMO resin was incubated with 1 μM Otulin (lanes 5 and 6), 1 μM Otulin plus 1 μΜ AMSH-LP (lanes 7 and 8), and Otulin plus AMSH-LP and then 0.5 M hydroxylamine (lanes 9 and 10), or 1 μM USP2 alone (lanes 11 and 12) (see Methods).

To establish whether these putative ester-bonds are catalysed by HOIL-1, we repeated the experiments with primary bone marrow-derived macrophages (BMDM) from mice expressing the E3 ligase-inactive HOIL-1[C458S] mutant. Similar to the RAW macrophage-like cell line, the ubiquitylated IRAK4 in primary BMDM from wild type (WT) mice is partially converted to deubiquitylated IRAK4 by incubation with hydroxylamine but the ubiquitylated IRAK4 formed in BMDM from HOIL-1[C458S] mice is resistant to this nucleophile (**Figs 1C and 1D**). The results establish that formation of the hydroxylamine-sensitive bonds in ubiquitylated IRAK4 are require HOIL-1 E3 ligase activity.

Treatment with Otulin, a deubiquitylase (DUB) that hydrolyses M1-Ub linkages specifically **(Keusekotten *et al*, 2013; Rivkin *et al*, 2013)**, reduced the size of the ubiquitin chains attached to IRAK4, without converting IRAK4 to either the mono-ubiquitylated or de-ubiquitylated species, while treatment with Otulin followed by treatment with AMSH-LP, a DUB that hydrolyses K63-Ub chains specifically **(Komander *et al*, 2009)**, largely converted ubiquitylated IRAK4 to monoubiquitylated and deubiquitylated species (**Figs 1E and 1F**). The observation that AMSH-LP causes partial cleavage of the first ubiquitin attached to some proteins has been noted previously **(Emmerich *et al***., **2013)**. Taken together, the results indicate that, similar to IRAK1 and IRAK2, the K63/M1-Ub chains attached to IRAK4 are initiated by both ester bonds and isopeptide bonds.

### Identification of serine, threonine, and lysine residues in myddosome components that undergo ubiquitylation during TLR7/8 signaling

Ubiquitin comprises 76 amino acid residues and terminates in the sequence Arg-Gly-Gly. To identify sites of ubiquitylation, a protocol has been developed in which ubiquitylated proteins are digested with trypsin to cleave the Arg-Gly bond between amino acid residues 74 and 75 of ubiquitin, generating isopeptides containing characteristic “Gly-Gly” (diGly) signatures that can be enriched by immunoprecipitation with an antibody recognising diGly-isopeptide sequences prior to identification of the peptides by mass spectrometry **(Xu *et al*, 2010)**. However, these antibodies do not recognise diGly sequences attached by ester bonds to the hydroxyl side chains of serine or threonine residues, and nor do they distinguish between lysine residues forming isopeptide bonds with ubiquitin or with the two ubiquitin-like modifiers (UBLs) ISG15 and NEDD8, both of which terminate in a diGly sequence. To circumvent these problems, proteins are therefore initially digested with LysC proteinase to generate ubiquitylated peptides in which amino acid residues 64-76 of ubiquitin are linked by isopeptide bonds to peptides from other proteins. These isopeptides are enriched by immunoprecipitation with “UbiSite”, an antibody that recognises amino acids 64-76 of ubiquitin but not the C-terminal peptides from other UBLs **(Akimov *et al*, 2018)**. Re-digestion with trypsin, now generates peptides in which the diGly signature from ubiquitin is linked to Ser/Thr or Lys residues, and which are then identified by mass spectrometry (MS).

The UbiSite method enabled us to identify sites of ester-linked ubiquitylation in IRAK4, IRAK2 and MyD88 (**Fig 2**) and sites of isopeptide-linked ubiquitylation in IRAK4, IRAK2, IRAK1 and MyD88 (**Figs EV1-3**). The ubiquitylation sites were only detected in R848-stimulated cells and not in unstimulated cells. The sites of ester-linked ubiquitylation were confined to the proline, serine, threonine-rich PST region of IRAKs 4 and 2, and the intermediate domain of MyD88 (**Fig 3**). Lys162 of IRAK2 is adjacent to Thr163, and Thr163 is located within the same tryptic peptide as Ser168, but no diubiquitylated peptides were detected in which both Thr163 and Ser168, Lys162 and Ser163, or Lys162 and Ser168 were ubiquitylated; nor was a peptide detected in which all three sites were ubiquitylated. These observations suggest that the ubiquitylation of these sites is mutually exclusive and consequently that each of these three sites of ubiquitylation are attached to different IRAK2 molecules. Sites of isopeptide-linked ubiquitylation were not only found in the PST and intermediate domains, but also in the kinase domain of IRAK4, the pseudokinase domain of IRAK2, the C-terminal domain of IRAK1 and the TIR (Toll-Interleukin Receptor) domain of MyD88 (**Fig 3)**. Together, these results establish that the ubiquitin chains attached to the components of myddosomes are indeed initiated by both ester and isopeptide bonds.

**Figure 2:**
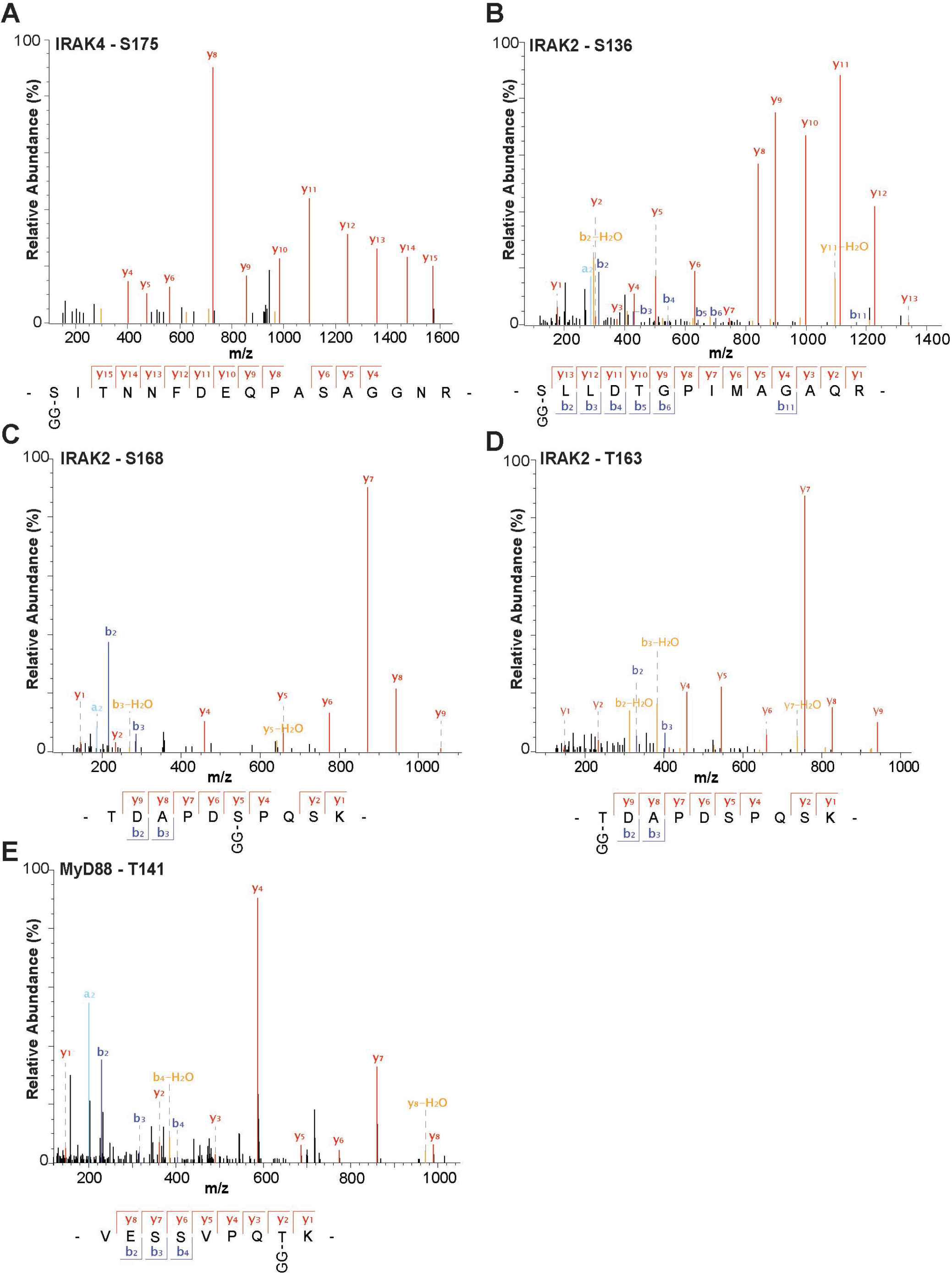
Ser and Threonine residues in Myddosome components shown to undergo R848-stimulated ubiquitylation in mouse RAW cells. A-E Mass fragmentation patterns of the indicated tryptic peptides from IRAK4 (A), IRAK2 (B-D), and MyD88 (E). “GG” represents the remnant of Ubiquitin attached to the specified amino acid in the target protein generated after tryptic digestion. Data information: The spectra shown were obtained 20 min after stimulation with R848.

**Figure 3:**
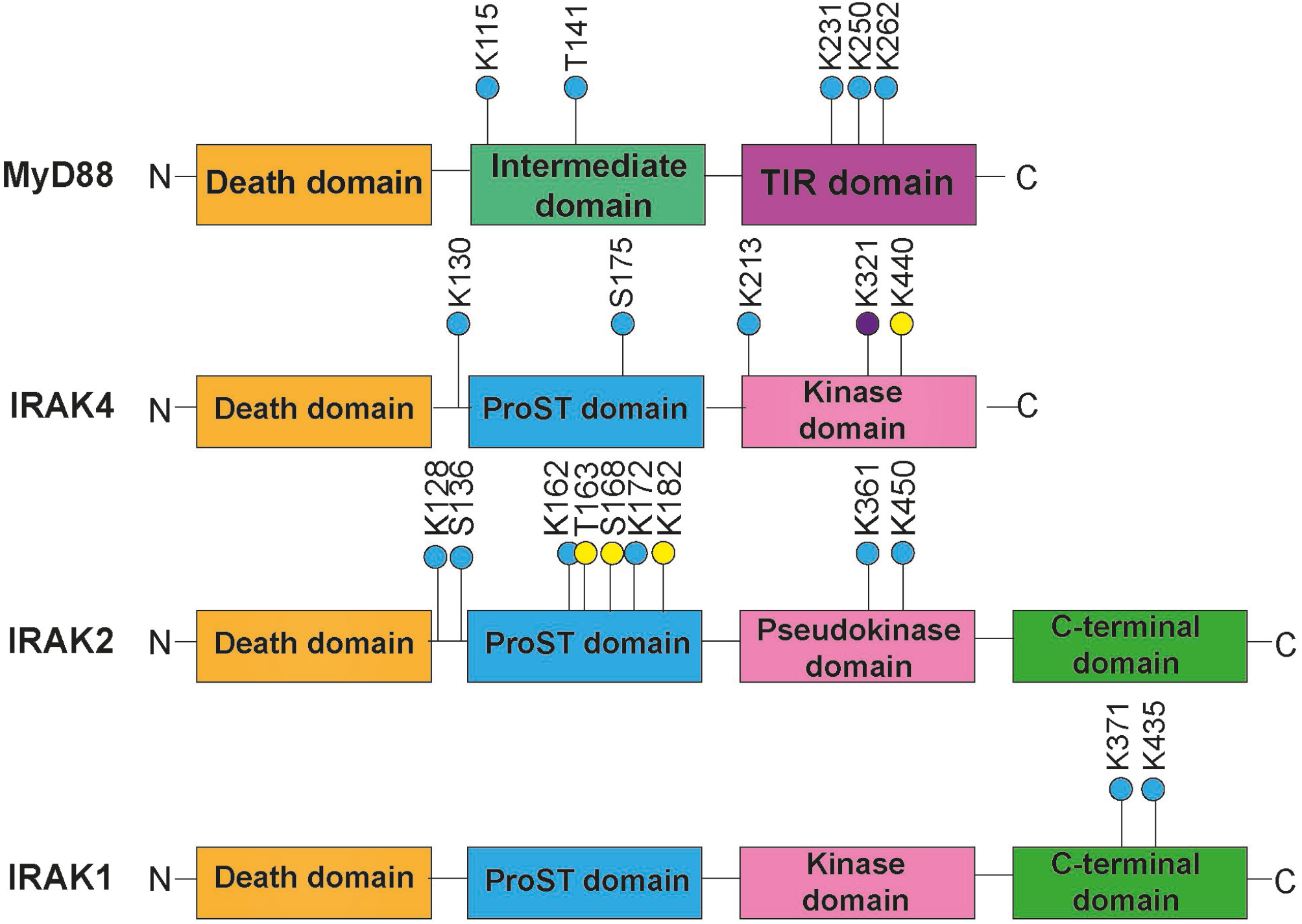
Location of ubiquitylated serine, threonine and lysine residues within the domain structures of IRAKs and MyD88. **(A)** The sites of ubiquitylation identified after stimulation for 20 min only (yellow circles), or at 6 h only (purple circles) or identified at both 20 min and 6h (blue circles). Abbreviations:-ProST domain, proline, serine, threonine-rich domain; TIR domain, Toll Interleukin-1 Receptor domain. The C-terminal domains of IRAK1 and IRAK2 contain TRAF6 interaction (Pro-Xaa-Glu) motifs where Xaa can be any amino acid.

We did not detect any sites of ester-linked ubiquitylation in IRAK1, which appear to be present based on the sensitivity of polyubiquitylated IRAK1 to cleavage with hydroxylamine (**Fig 1A**) **(Kelsall *et al***., **2019)**. This may be explained if, like IRAK2 and IRAK4, these ester bonds are located within the PST domain, because the PST domain of mouse IRAK1 only contains two lysine (Lys101 and Lys147) and two arginine residues (Arg 100 and Arg160). Since the ubiquitylation of a lysine residue is known to prevent tryptic cleavage, the expected tryptic peptides derived from the PST region of IRAK1 would comprise amino acid residues 101-160 or 63-160, if Lys101 and Lys147 were ubiquitylated and depending on whether the ubiquitylation of Lys101 also prevents tryptic cleavage at Arg 100. Peptides of this size can be difficult to detect by mass spectrometry and suggests that digestion with additional proteases will be needed to generate smaller ubiquitylated peptides that can be identified more easily. Lys134 and Lys180 of human IRAK1, the residues equivalent to Lys101 and Lys147 of mouse IRAK1, are reported to be major and minor sites of isopeptide ubiquitylation, based on overexpression studies in HEK293 cells in which either or both of these lysine residues were mutated to arginine (Conze *et al*, 2008).

### Ester-linkages are formed between two ubiquitin molecules

The only other protein that we detected in the Halo-NEMO pull-downs that was linked to ubiquitin by an ester bond was ubiquitin itself. Remarkably, we found that four different Ser/Thr residues in ubiquitin formed ester bonds with the C-terminus of another ubiquitin molecule namely Thr12 (**Fig 4A**), Thr14 (**Fig 4B**), Ser20 (**Fig 4C**) and Thr22 (**Fig 4D**). The linkages between Thr12, Ser20 and Thr22 were detected after stimulation for 20 min with R848 and the R848-dependent ubiquitylation of Thr12 was also detected after stimulation for 6 h. **(Fig 3)**. The ubiquitylation of Thr14 was also identified in unstimulated cells.

**Figure 4:**
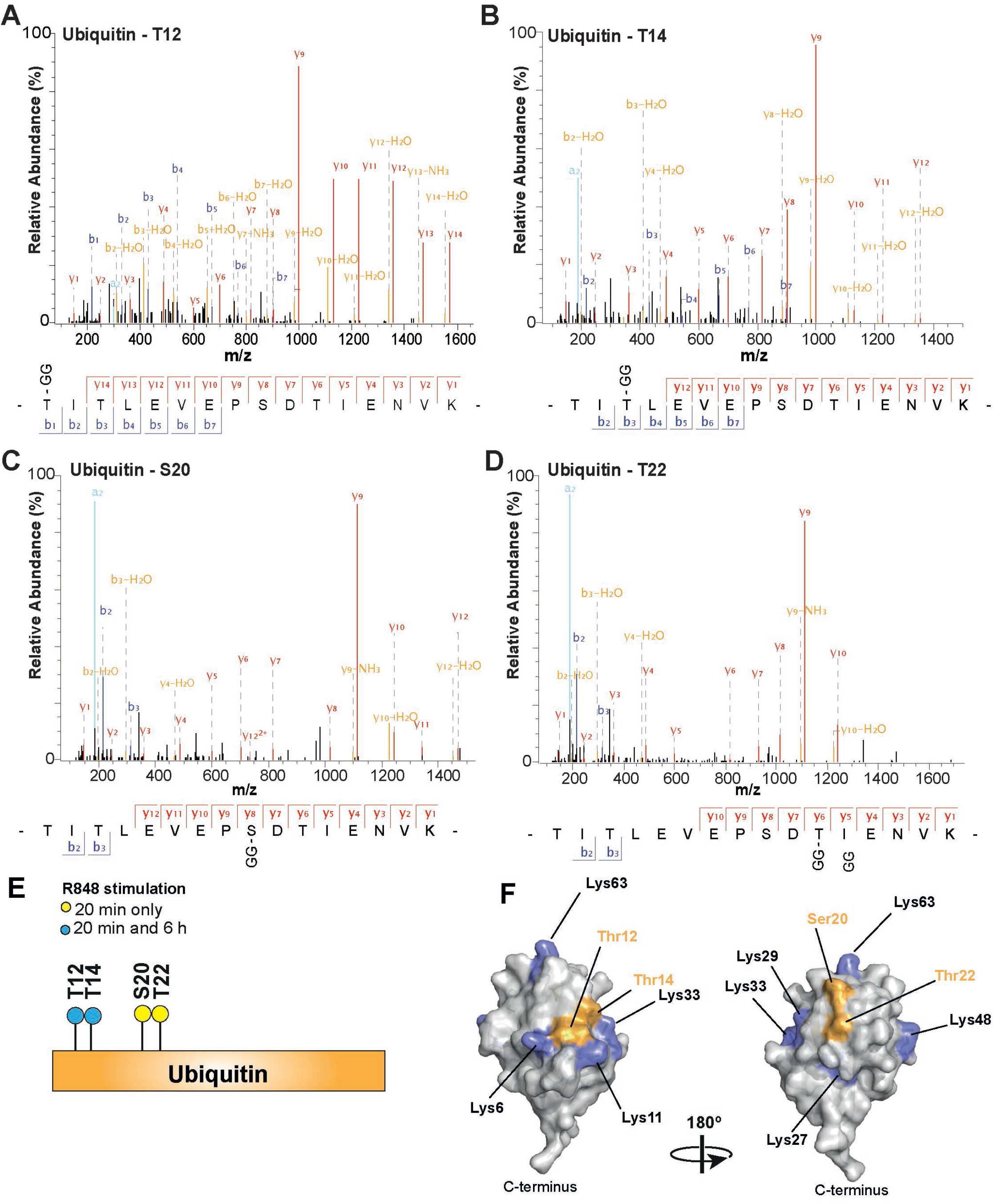
Ester-linked ubiquitin dimers formed in R848-stimulated RAW cells. **A-D** Mass fragmentation patterns of tryptic peptides comprising amino acid residues 12-27 of ubiquitin, which were formed in R848-stimulated RAW cells and captured on Halo-NEMO beads. Each of the four peptides contains a “GG” signature and their fragmentation patterns show that they are ubiquitylated at Thr12, Thr14, Ser 20 or Thr22. **E** Schematic showing the time at which each ubiquitin dimer was identified after stimulation with R848. All four ester-linked ubiquitylation sites were detected after stimulation for 20 min with R848 (yellow circles) but Thr12 and Thr14 were the only ubiquitylation sites detected after stimulation for 6 h (blue circles). **F** The three-dimensional structure of ubiquitin (PDB: 1UBQ) showing the location of Thr12, Thr14, Ser20 and Thr22 and the seven lysine residues **(Vijay-Kumar *et al***., **1987)**. Data information: The spectra shown were obtained 20 min after stimulation with R848.

Until now, eight types of ubiquitin dimer have been identified in cells in which the C-terminus of one ubiquitin can be linked to the ε-amino groups of any of the seven lysine residues or the α-amino group of the N-terminal methionine of another ubiquitin. The present study therefore increases the number ubiquitin linkage types that are formed in cells from 8 to 12. HOIL-1 is probably the E3 ligase that ubiquitylates Thr12, Ser20 and Thr22 because it has been shown to form these three ester-linked ubiquitin dimers *in vitro* (**Kelsall *et al***., **2019**). Whether HOIL-1 is also the E3 ligase that ubiquitylated Thr14 of ubiquitin will require further experimentation. The LUBAC-catalysed formation of an ester-bond between Thr55 of ubiquitin and the C-terminal carboxylate of another ubiquitin molecule *in vitro* has also been reported **(Rodriguez Carvajal *et al***., **2021)**. We did not detect ester bonds linking Thr55 of ubiquitin to another ubiquitin molecule in our experiments, but this does not exclude the possibility that this dimer is formed in response to other stimuli and/or in other cells. Although Thr12, Thr14, Ser 20 and Thr22 are all located in the same tryptic peptide comprising amino acid residues 12-27 of ubiquitin, we did not detect any peptide ubiquitylated at more than one of these sites. Thus, four separate ester-linked ubiquitin dimers are formed in response to R848.

It is a paradigm that different ubiquitin dimers adopt distinct conformations that are recognised by specific ubiquitin-binding proteins, which function to decode the ubiquitin signal [reviewed **(Radley *et al*, 2019)**]. Thr12/Thr14 and Ser20/Thr22 are situated on the surface of ubiquitin but on opposite sides of the molecule (**Fig 4F**). This suggests that the structures of these two pairs of ubiquitin dimers are likely to differ markedly and, consequently, may interact with different proteins. It will clearly be important to determine the structures of each ester-linked ubiquitin dimer by X-ray crystallography and to identify their binding partners in cells. The binding partners may include proteins that stimulate or suppress innate immune signaling, explaining why humans with HOIL-1-deficiency display a combination of immunodeficiency and auto-inflammation **(Boisson *et al***., **2012; Phadke *et al***., **2020)**.

The ester-linked ubiquitin dimers were identified after capturing them on Halo-NEMO resin. Since NEMO interacts with M1-Ub dimers, this suggests that ester-linked ubiquitin dimers may be attached covalently either to M1-Ub oligomers and/or to the K63/M1-Ub hybrids captured by the Halo-NEMO resin **(Emmerich *et al***., **2013)**. The high molecular mass ubiquitin chains formed by stimulating mouse embryonic fibroblasts with tumour necrosis factor (TNF), which were detected by immunoblotting with an M1-Ub-specific antibody, also contain ester linkages as judged by their reduced size after incubation with hydroxylamine **(Rodriguez Carvajal *et al***., **2021)**. However, whether the ester linked ubiquitin dimers are linked to M1-Ub and/or K63-Ub linkages or to each other, or even to other proteins captured by the Halo-NEMO resin that we did not detect in our study, requires more detailed analysis of the composition and topology of the ubiquitin chains formed in TLR-stimulated cells.

The identities of the DUB(s) that deubiquitylate(s) the ester-linked ubiquitin bonds that we have detected in macrophages is another interesting topic for future research. Several DUBs of the USP (ubiquitin-specific protease) family, the Machado-Joseph disease (MJD) family, and vOTU, a virally encoded member of the OTU (Ovarian Tumour) family of DUBs, have been shown to hydrolyse a ubiquitylated threonine substrate linked by an ester-bond *in vitro* **(De Cesare *et al*, 2021)**. However, USP family members and vOTU also hydrolyse isopeptide bonds, indicating that ester-linked and isopeptide-linked ubiquitin dimers may not necessarily be deubiquitylated by different DUBs.

Interestingly, Ser136 of IRAK2, one of the sites of ubiquitylation identified in this study, has been reported to undergo phosphorylation when mouse BMDM are stimulated with lipopolysaccharide, an activator of TLR4 (Weintz *et al*, 2010). Moreover, the phosphorylation of Thr12 (Walser *et al*, 2020), Thr14 (Zhou *et al*, 2013), Ser20 **(Lundby *et al*, 2012)** and Thr22 (Swaney *et al*, 2013) of ubiquitin have all been reported in a variety of mammalian cells. Clearly, one potential role for the phosphorylation of these sites could be to prevent their ubiquitylation but, conversely, ubiquitylation may prevent phosphorylation. Since ubiquitylation and phosphorylation of the same amino acid residue is mutually exclusive, this could be an interesting way in which protein kinases regulate the actions of E3 ligases forming ester bonds and vice versa.

## Materials and Methods

### Plasmids and proteins

The plasmids utilised in this study and glutathione-S-Transferase (GST) fusions of human GST-Otulin (DU43487), human AMSH-LP[264–436] (DU15780) and GST-USP2 (DU13025) were produced by MRC Reagents and Services and can be requested at http://mrcppureagents@dundee.ac.uk. The preparation of Halo-tagged NEMO and its covalent attachment to Halo-Link Resin (Promega) have been described (Emmerich *et al*., 2013). LysC proteinase was from Thermo Fischer Scientific (#90051), trypsin from Promega (#V511A) and the protein phosphatase from bacteriophage λgt10 (phage λ phosphatase) (Cohen & Cohen, 1989) from New England Biolabs (#P0753S).

### Antibodies

The antibodies recognising IRAK1 (Cell Signalling Technology #4504S) and IRAK2 (Abcam #ab62419) were purchased from the sources indicated in parentheses. The antibody recognising IRAK4 phosphorylated at both Thr345 and Ser346 was a gift from Dr Vikram Rao (Pfizer, Boston, MA, USA). This antibody was used in this study because it is more sensitive than any commercially available IRAK4 antibody that we have tested. The antibodies specific for Lys63-linked and Met1-linked ubiquitin oligomers were gifts from Dr Vishva Dixit (Genentech, San Francisco, USA).

### Cell culture, stimulation and lysis

RAW264.7 cells (RAW cells) were maintained in Dulbecco’s modified Eagle’s Medium (DMEM) supplemented with 10% (v/v) foetal bovine serum, 2 mM L-glutamine, 100 Units/ml penicillin and 0.1 mg/ml streptomycin. Bone marrow from HOIL-1[C458S] mice **(Kelsall *et al***., **2019)** and wild type littermates was differentiated into primary bone marrow-derived macrophages using L929 preconditioned medium **(Pauls *et al*, 2013)**. Cells were stimulated with or without 250 ng/ml R848, washed rapidly with ice-cold PBS and lysed in 50 mM Tris-HCl pH 7.5, 1 mM EGTA, 1 mM EDTA, 1% (v/v) Triton X-100, 270 mM sucrose, containing the protein phosphatase inhibitors 10 mM glycerol-2-phosphate, 50 mM sodium fluoride, 5 mM sodium pyrophosphate, 1 mM sodium orthovanadate, and the proteinase inhibitors 1 mM phenylmethanesulphonyl fluoride, cOmplete protease inhibitor (one tablet per 50 ml) (Roche) and 100 mM iodoacetamide to inhibit deubiquitylase activities. Cell lysates were centrifuged at 17,000 x g for 15 min at 4°C and the supernatant, termed cell extract was removed and used for experimentation. Protein concentrations in cell extracts were measured by the Bradford Assay **(Bradford, 1976)**.

### Capture of proteins on Halo-NEMO beads and treatment with deubiquitylases and hydroxylamine

Halo-NEMO resin **(Emmerich *et al***., **2013)** was washed twice with 6M urea solution and five times with 50 mM Tris/HCl pH 7.5, 1% (v/v) Triton X-100. Following incubation overnight with cell extract (2 mg per 20 μl packed Halo-NEMO resin), the resin was collected by centrifugation, washed twice with 50 mM Tris/HCl pH 7.5, 500 mM NaCl, 1% (v/v) Triton X-100 and twice with 50 mM Tris/HCl pH 7.5, 1% (v/v) Triton X-100 and the supernatant discarded.

The packed beads were resuspended in 50 mM HEPES pH 7.5, 100 mM NaCl, 2 mM DTT, 0.01% (w/v) Brij-35 and incubated for 1 h at 37 °C without or with one more of the deubiquitylases Otulin, AMSH-LP and USP2 (each at 1 μM) as indicated in the figure legends. The beads were washed twice with 50 mM Tris/HCl pH 7.5, 1% (v/v) Triton X-100, once with 50 mM Tris/HCl pH 7.5 and resuspended in 19 mM sodium carbonate, 22 mM sodium bicarbonate pH 9.0 and incubated for 1 h at 37 °C with or without 0.5 M hydroxylamine. The beads were washed twice with 50 mM Tris/HCl pH 7.5, 1% (v/v) Triton X-100 and proteins eluted by resuspension for 10 min at 37 °C in LDS sample buffer (Invitrogen) diluted four-fold in 2.5% (v/v) 2-mercaptoethanol. The eluted proteins were passed through a Spin-X centrifuge filter (Corning Costar) and subjected to SDS-PAGE using precast 4-12% gels and transferred to PVDF membranes prior to immunoblotting.

### Mapping of ubiquitylated amino acid residues using the UbiSite antibody

250 mg cell extract protein was incubated with 2 ml packed Halo-NEMO beads. After completion of the washing steps, the beads were resuspended in 50 mM HEPES pH 7.5, 100 mM NaCl, 2 mM DTT, 0.01% (w/v) Brij-35 and incubated for 30 min at 37 °C with 1000 U of phage λ phosphatase. The beads were washed twice with 50 mM Tris/HCl pH 7.5, 1% (v/v) Triton X-100, five times with 50 mM Tris/HCl pH 7.5, and then incubated for 5 min at 37 °C with 5% (w/v) Rapigest. After diluting 5-fold with 50 mM Tris/HCl pH 7.5 and incubation for 10 min at 37 °C, the eluted proteins were separated from the beads using a Spin-X centrifuge filter. The flowthrough containing the eluted proteins was made 2 mM in DTT, incubated for 30 min at room temperature and then alkylated for a further 30 min with 11 mM chloroacetamide (CAA). The samples were diluted 10-fold with 50 mM Tris/HCl pH 7.5 to reduce the Rapigest concentration to 0.1%, then digested overnight at 37 °C with LysC proteinase at a concentration of 1:100 (w/w) enzyme to protein. Digestion was terminated by the addition of trifluoroacetic acid (TFA) to a final concentration of 2% (v/v) and incubated for 1 h at 37 °C. Following centrifugation for 10 min at 10,000 x g, the supernatant was removed and desalted using a WATERS C_18_ Cartridge (WAT054955) according to the manufacturer’s instructions. The eluted peptides were lyophilised, reconstituted in 50 mM MOPS pH 7.2, 10 mM Na_2_HPO_4_, 50 mM NaCl, enriched by immunoprecipitation with the Ubisite antibody and analysed as described before (Akimov *et al*., 2018).

Mass spectrometric analyses were performed essentially as described **(Trulsson *et al*, 2022)**, with a few modifications. An Orbitrap Exploris 480 mass spectrometer (ThermoFisher Scientific) was used in a data-dependent mode (DDA method) by shifting from full-scan event to the top 12 MS/MS scans. The instrument was operated in positive polarity mode with the following parameters for full-scan acquisitions: the normalized automatic gain control (AGC) target value was set to 300%, resolution was 60,000 with a scan range of 350-1500 m/z and a maximum ion injection time (IT) of 25 ms. The ion charge range was set from 2 to 6 charges. MS/MS fragmentation of precursors were obtained by HCD (higher-energy collisional dissociation) with a normalized collisional energy of 30. Resolution of MS/MS scans was 30,000, maximum IT of 54 ms and the isolation window was 1.2 m/z. The dynamic exclusion window was set to 60 se to prevent the same peptide sequences from repeating. Raw MS files were processed through Proteome Discoverer software (P.D. version 2.5.0.400; ThermoFisher Scientific) using the Sequest HT search engine with the following parameters: the precursor mass tolerance was set to 10 ppm and the fragment mass tolerance was 0.02 Da, the protease trypsin was allowed a maximum of two missed cleavage sites and a minimum peptide length of six amino acid residues. Carbamidomethylation (+57.021 Da) of Cysteine residues was set as fixed modification. Oxidation of Methionine (+15.995 Da) and diGly modification on Lysine, Serine, and Threonine (+114.043 Da) were set as variable modifications. Data were searched against the Uniprot Mouse database (from April 2022) using Percolator node of P.D. software to estimate FDR (less than 1%). All spectra with diGly modifications on serine and threonine residues presented in the manuscript were verified manually.

## Data Availability

All data mentioned in this paper have been provided in the main article.

### Acknowledgements

We thank our colleagues Ian Kelsall and Ron Hay for their helpful suggestions. This study was supported by Wellcome Trust Investigator Award 209380/Z/17/Z (to PC) and by a Wellcome Trust Prize Studentship (to EM). The work in the BB group was supported in part by the Danish National Research Foundation (DNRF grant no. 141, ATLAS), the Independent Research Fund Denmark (DFF—8022-00051) and the INTEGRA research infrastructure (NNF20OC0061575) fund of the Novo Nordisk Foundation.

## Author Contributions

EM: Investigation, formal analysis, methodology, writing. VA: acquisition and formal analysis of mass spectrometry data, investigation, methodology. BB: acquisition and formal analysis of mass spectrometry data, investigation, methodology, supervision, funding acquisition. PC: conceptualization, formal analysis, supervision, funding acquisition, writing.

## Expanded View Figure Legends

**Fig EV1.**
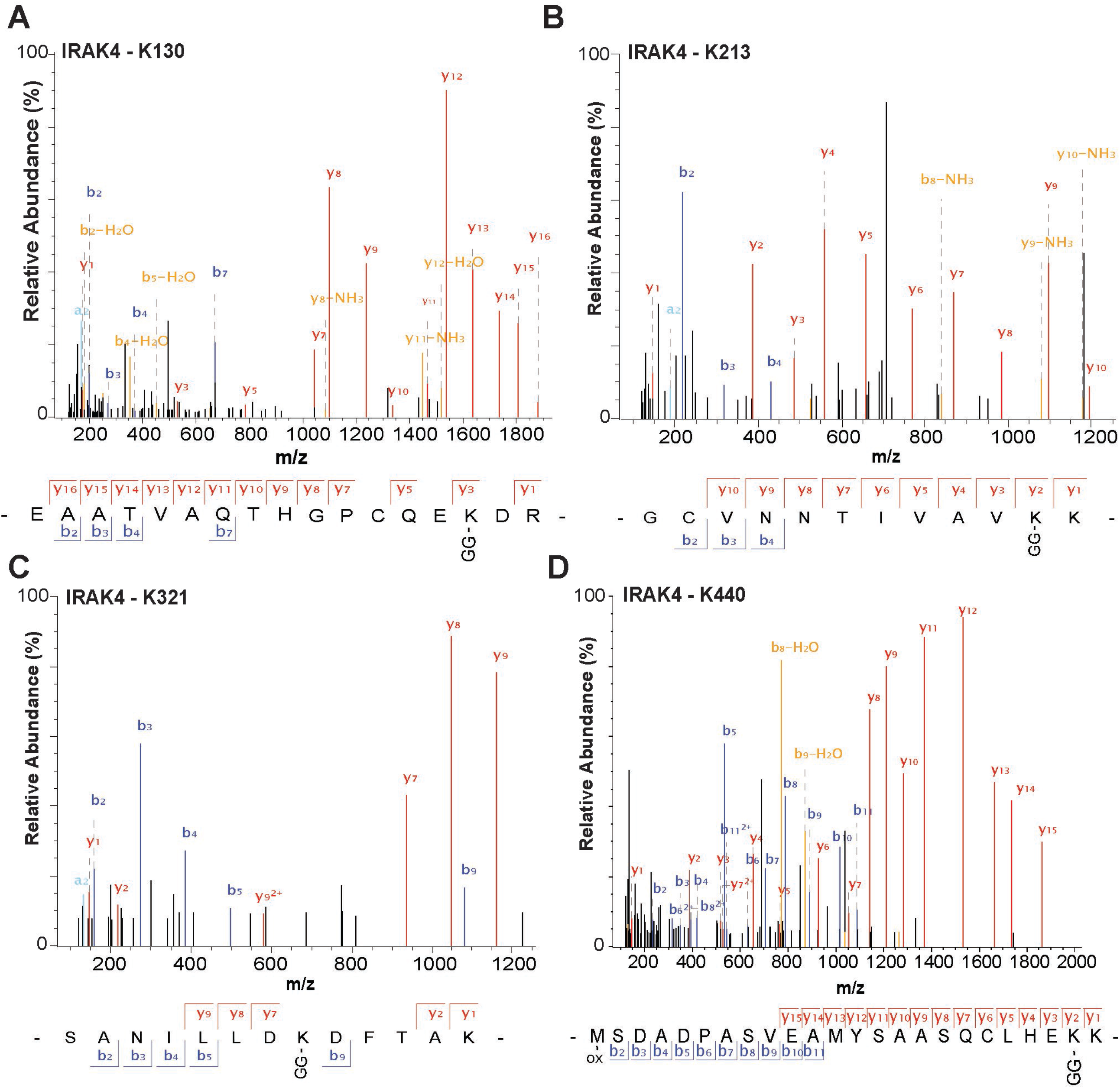
Isopeptide-linked ubiquitylation sites in IRAK4 formed in response to R848 in mouse RAW cells. Mass fragmentation patterns of the indicated tryptic peptides from IRAK4. “GG” represents the remnant of Ubiquitin attached to the specified amino acid in the target protein generated after trypsin digestion. Data information: The spectra shown were obtained 20 min after stimulation with R848 except for the peptide containing Lys321 which was obtained after stimulation for 6. h.

**Fig EV2.**
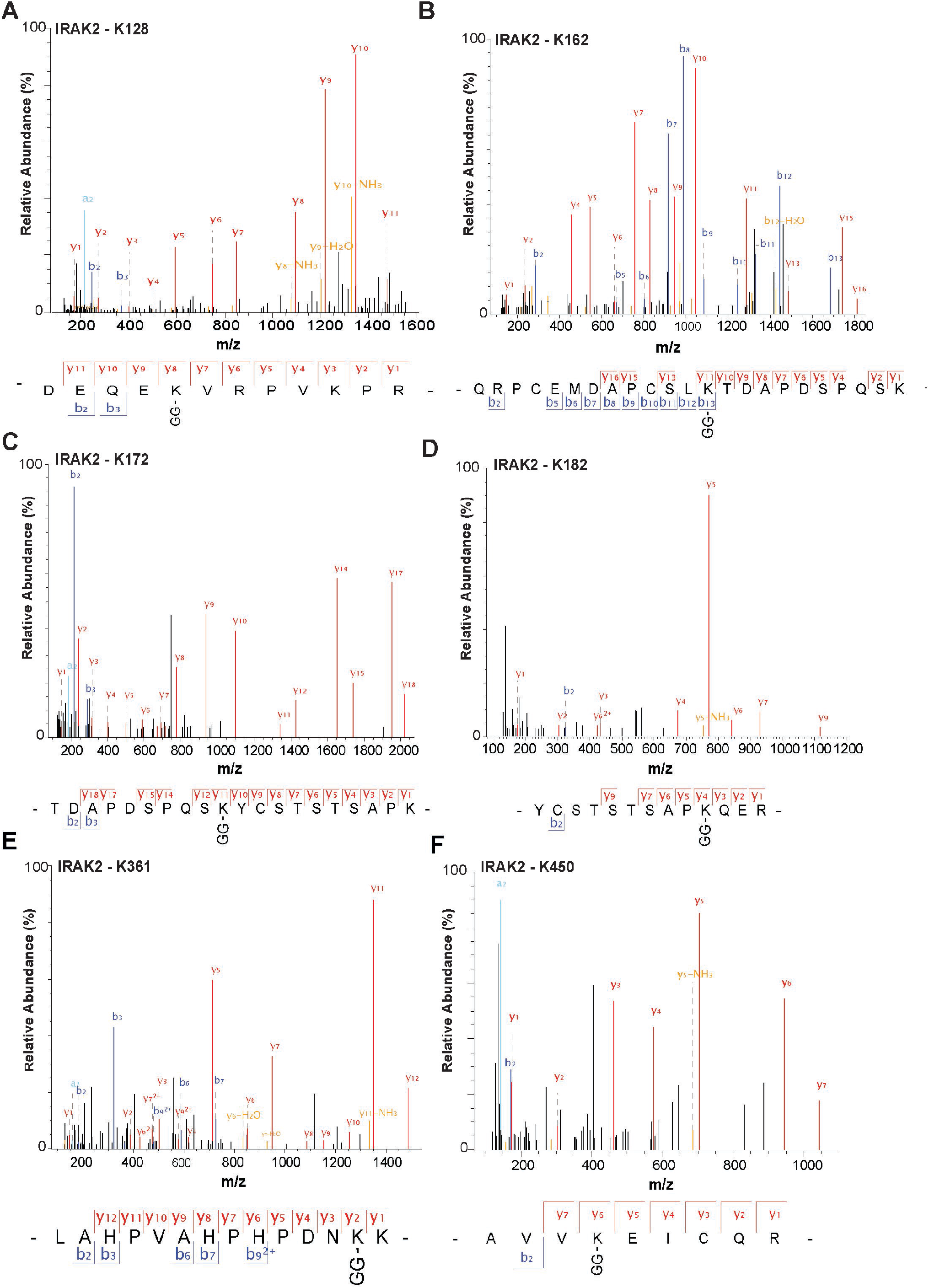
Isopeptide-linked ubiquitylation sites in IRAK2 formed in response to R848 in mouse RAW cells. Mass fragmentation patterns of the indicated tryptic peptides from IRAK2. “GG” represents the remnant of Ubiquitin attached to the specified amino acid in the target protein generated after trypsin digestion. Data information: The spectra shown were obtained 20 min after stimulation with R848.

**Fig EV3.**
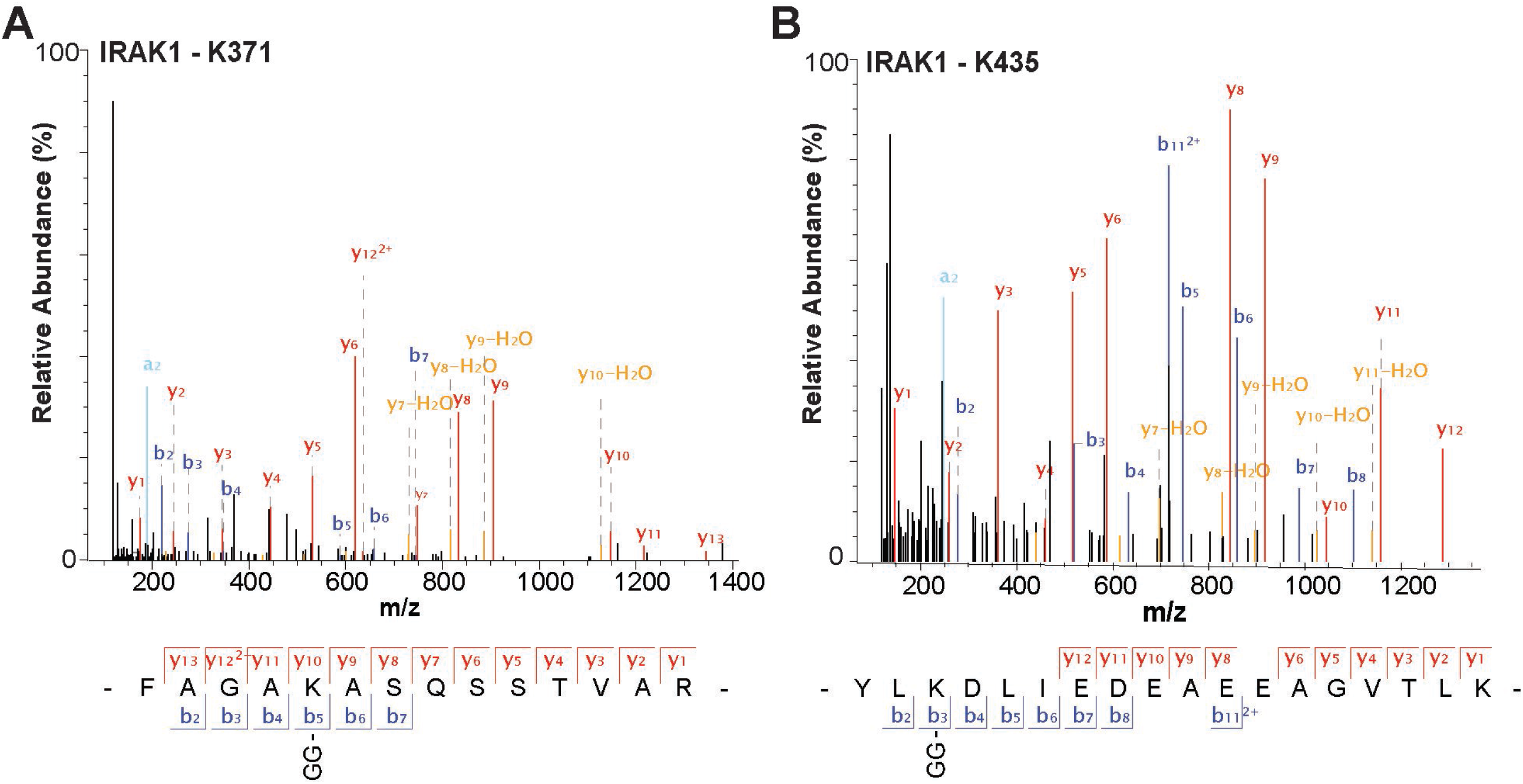
Isopeptide-linked ubiquitylation sites in IRAK1 formed in response to R848 in mouse RAW cells. Mass fragmentation patterns of the indicated tryptic peptides from IRAK1. “GG” represents the remnant of Ubiquitin attached to the specified amino acid in the target protein generated after trypsin digestion. Data information: The spectra shown were obtained 20 min after stimulation with R848

**Fig EV4.**
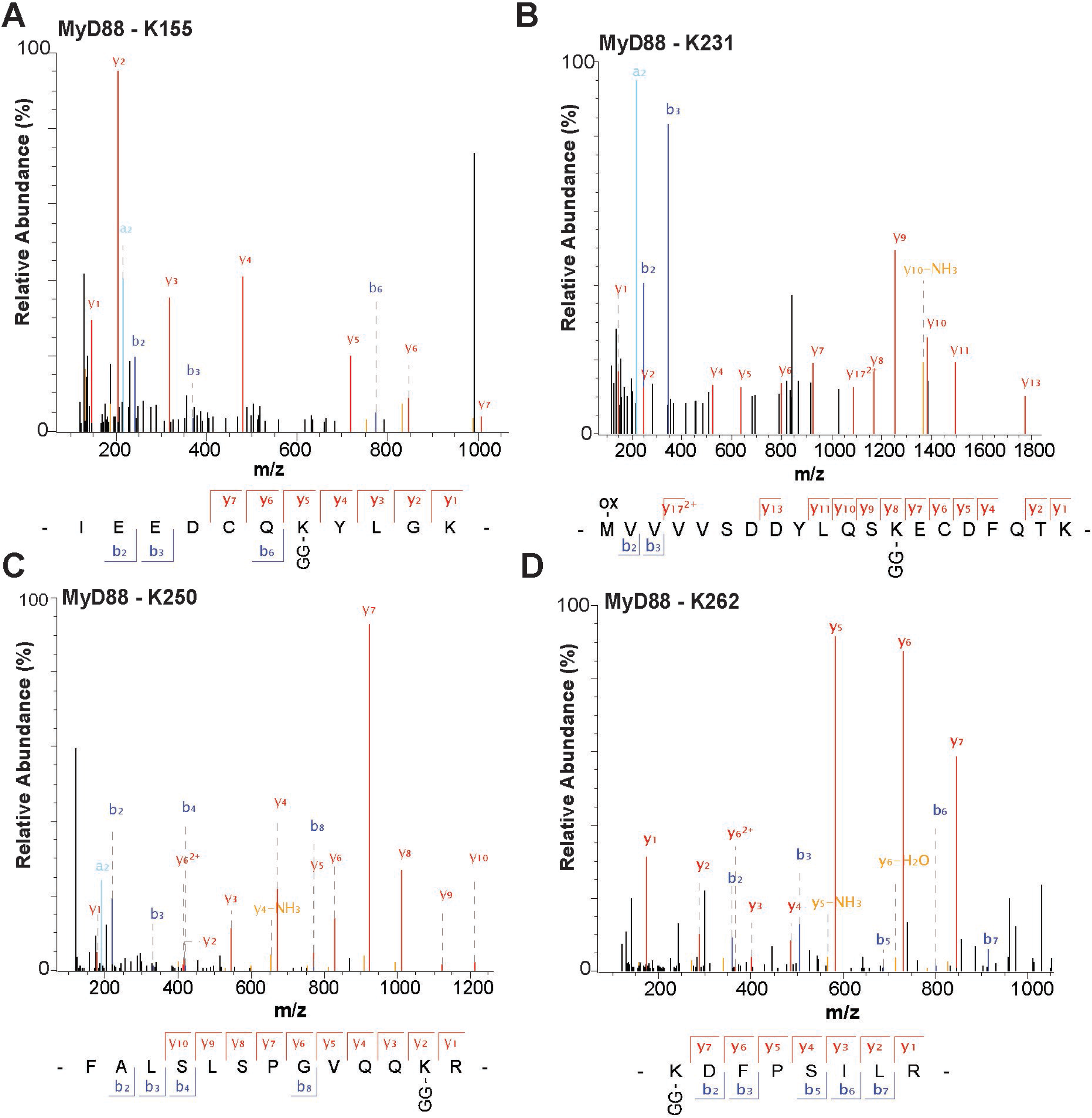
Isopeptide-linked ubiquitylation sites in MyD88 formed in response to R848 in mouse RAW cells. Mass fragmentation patterns of the indicated tryptic peptides from MyD88. “GG” represents the remnant of Ubiquitin attached to the specified amino acid in the target protein generated after trypsin digestion. Data information: the spectra shown were obtained 20 min after stimulation with R848.

